# Visual experience has opposing influences on the quality of stimulus representation in adult primary visual cortex

**DOI:** 10.1101/2022.05.17.492357

**Authors:** Brian B. Jeon, Thomas Fuchs, Steven M. Chase, Sandra J. Kuhlman

**Affiliations:** Department of Biomedical Engineering, Carnegie Mellon University; Center for the Neural Basis of Cognition, Carnegie Mellon University; Neuroscience Institute, Carnegie Mellon University; Department of Biological Sciences, Carnegie Mellon University

## Abstract

Transient dark exposure, typically 7-10 days in duration, followed by light reintroduction is an emerging treatment for improving the restoration of vison in amblyopic subjects whose occlusion is removed in adulthood. Dark exposure initiates homeostatic mechanisms that together with light-induced changes in cellular signaling pathways result in the re-engagement of juvenile-like plasticity in the adult such that previously deprived inputs can gain cortical territory. It is possible that dark exposure itself degrades visual responses, and this could place constraints on the optimal duration of dark exposure treatment. To determine whether eight days of dark exposure has a lasting negative impact on responses to classic grating stimuli, neural activity was recorded before and after dark exposure in awake head-fixed mice using 2-photon calcium imaging. Neural discriminability, assessed using classifiers, was transiently reduced following dark exposure; a decrease in response reliability across a broad range of spatial frequencies accounted for the disruption. Both discriminability and reliability recovered. Fixed classifiers were used to demonstrated that stimulus representation rebounded to the original, pre-deprivation state, thus DE did not appear to have a lasting negative impact on visual processing. Unexpectedly, we found that dark exposure significantly stabilized orientation preference and signal correlation. Our results reveal that natural vision exerts a disrupting influence on the stability of stimulus preference for classic grating stimuli, and at the same time improves neural discriminability for both low and high spatial frequency stimuli.

## Introduction

Sensory cortex is highly malleable early in life. During postnatal development, cortical territory rapidly expands and contracts to represent active and inactive inputs, respectively. These large-scale changes are mediated by age-restricted experience-dependent synaptic refinement at the level of individual postsynaptic neurons^1^. Primary visual cortex (V1) is particularly sensitive to visual deprivation during this period. Classic experiments demonstrate that in order for sensory pathways to drive cortical responses in the adult, continuous binocular input is required during postnatal development^2,3^. Experimentally induced monocular deprivation in young animals results in a rapid decrease in the strength of deprived inputs and with a delay, the response of intact inputs is potentiated. Depression of deprived inputs occurs at a synaptic level and outlasts the perturbation. On the other hand, the same peripheral perturbation in adults does not induce a rapid decrease in response strength, although potentiation of the intact input does proceed^4,5^. Furthermore, in the young, the impact of deprivation on excitatory neuron activity levels is compensated for by a decrease in evoked inhibitory neuron activity. This form of compensation is a direct consequence of deprivation and does not occur in the adult^6^. Delineating the molecular basis and cellular signaling pathways responsible for restricting plasticity in the adult is an active area of investigation. For example, consistent with the above observations, extracellular matrix perineuronal nets (PNNs) which preferentially surround parvalbumin inhibitory neurons and create a barrier for synaptic remodeling, are resistant to degradation in the adult. Interventions that target the break-down of PNNs in adult V1 are effective in restoring juvenile-like plasticity and allow previously unused inputs to regain the ability to drive cortical neurons^2,7–11^. Taken together, there is a general consensus that in the adult, the inputs that are established during postnatal development retain a limited amount of plasticity throughout life. However, in contrast to the young, established inputs are not lost following disuse, and in the absence of additional training or treatment, new input patterns from previously occluded sources are not readily integrated into the existing networks of adults^3^. As such, restoring vision to subjects that have matured without binocular input during early postnatal development is a recognized challenge^3,12,13^.

To successfully restore vision, the newly opened inputs must compete with non-deprived eye inputs to drive cortical responses. An emerging treatment for animals with experimentally induced amblyopia, across a diverse range of species, is to transiently expose subjects to darkness, followed by light reintroduction^14–21^. Accumulating evidence indicates that dark exposure (DE) followed by light reintroduction (LRx) re-activates juvenile plasticity and is sufficient to restore deprived-eye input responses to grating stimuli. Notably, light reintroduction effectively degrades PNNs^17^. Thus, this critical form of juvenile plasticity is reactivated and likely contributes to the effectiveness of DE in restoring responsiveness of the deprived pathway^7^.

Ideally treatments such as DE followed by LRx would not have a negative impact on on-going visual processing carried out by the intact pathway. In other words, effective treatments would facilitate the integration of new information without perturbing existing functionality. Based on the studies cited above, it is not expected that in the adult basic responsiveness to visual stimuli following transient dark exposure would be lost. However, previous work demonstrated that closing one eye is sufficient to transiently disrupt stimulus representation, in the adult. Although individual neurons remain responsive, the pattern of activity evoked in V1 is disturbed, including orientation preference and pairwise signal correlation among simultaneously recorded neurons^22^. Similarly, although mean firing rates are largely similar across daily light-dark transitions, pairwise correlations are significantly stronger during vision^23^. This raises the possibility that deprivation such as DE could increase the rate of representational drift^24^, placing a burden on downstream areas to update the manner in which information is readout. Indeed, in association cortex, continuous experience is required for drift to remain low^25^. Given recent evidence that V1 exhibits characteristics similar to downstream cortical areas in the visual hierarchy^24^, higher-order relationships among neurons, beyond the basic responsiveness of individual neurons^26^, could be altered by the interruption of continuous visual input.

To determine whether transient DE disrupts or otherwise influences stimulus representation beyond basic responsiveness, neural responses to grating stimuli that covered a broad range of spatial frequencies were recorded using 2-photon calcium imaging in awake head-fixed mice before and after transient dark exposure. Calcium imaging has the advantage that individual neurons can be readily longitudinally tracked^27^. Tuning stability, and response reliability were assessed before and after DE. In addition, neural discriminability and the rate of representational drift were quantified using classifiers. We found that in contrast to monocular deprivation^22^, DE did not degrade pairwise signal correlation when immediately assessed after DE, and in fact stabilized orientation preference in neurons that remained tuned. Similarly, light reintroduction did not degrade pairwise signal correlation of tuned neurons. These data indicate that changes in signal correlation induced by monocular deprivation in adulthood are likely a result of imbalanced input, rather than reduced drive from the periphery.

However, similar to association areas, an increase in the rate of representation drift was detected when visual input was interrupted by DE. A decrease in the trial-to-trial reliability of stimulus responsiveness accounted for the transient change. Reliability was restored within 8 days of light reintroduction, and the representation rebounded to its original form. Thus when used for the treatment of amblyopia, neither DE or LRx are expected to have a persistent negative impact on existing visual processing, although the effectiveness of perceptual training^28^ immediately following DE may be influenced by a transient increase in representational drift. Furthermore, our results establish that although natural vision has a disrupting influence on tuning stability to simple grating stimuli, exposure to naturalistic statistics in the home-cage environment improves neural discriminability in the adult.

## Results and Discussion

To assess the impact of dark exposure (DE) on tuning stability and neural discriminability, the activity of individual layer 2/3 excitatory V1 neurons was imaged in response to randomized presentations of static grating stimuli using 2-photon microscopy in head-fixed transgenic mice positioned atop a floating spherical treadmill, expressing the calcium indicator GCaMP6f driven by the EMX1 promoter. Our goal was to assess the stability of stimulus representation before and after DE. Therefore prior to DE, 2 baseline imaging sessions were recorded, referred to here as Baseline 1 (B1) and Baseline 2 (B2), respectively. The acquisition of two baseline sessions allowed the stability of tuning to grating stimuli of varying orientation and spatial frequency to be assessed before DE was initiated. A third imaging session was recorded after DE, and is referred to as the Post-DE (pDE) session. A final forth imaging session was acquired after 8±1 days of light reintroduction (LRx), referred to as the Recovery (Rec) session. These 4 sessions were used to define 3 experimental conditions: Control, DE, and LRx (**Fig. 1 A,B**). A total of six mice were included in the study (see **Table 1** for sex and age information). Locomotion and pupil diameter were monitored; trials in which locomotion or eye blinking were detected were removed from analysis.

**Table 1.** Animal information.

| Animal ID | Animal label | Sex | Age on the first imaging session (days) |
| --- | --- | --- | --- |
| 2452_1R | mouse 1 | M | 83 |
| 2452_1R1L | mouse 2 | M | 83 |
| 2454_1R | mouse 3 | M | 86 |
| 2472_1L | mouse 4 | F | 59 |
| 2473_1R | mouse 5 | M | 45 |
| 2474_1R1L | mouse 6 | F | 56 |

**Figure 1.**
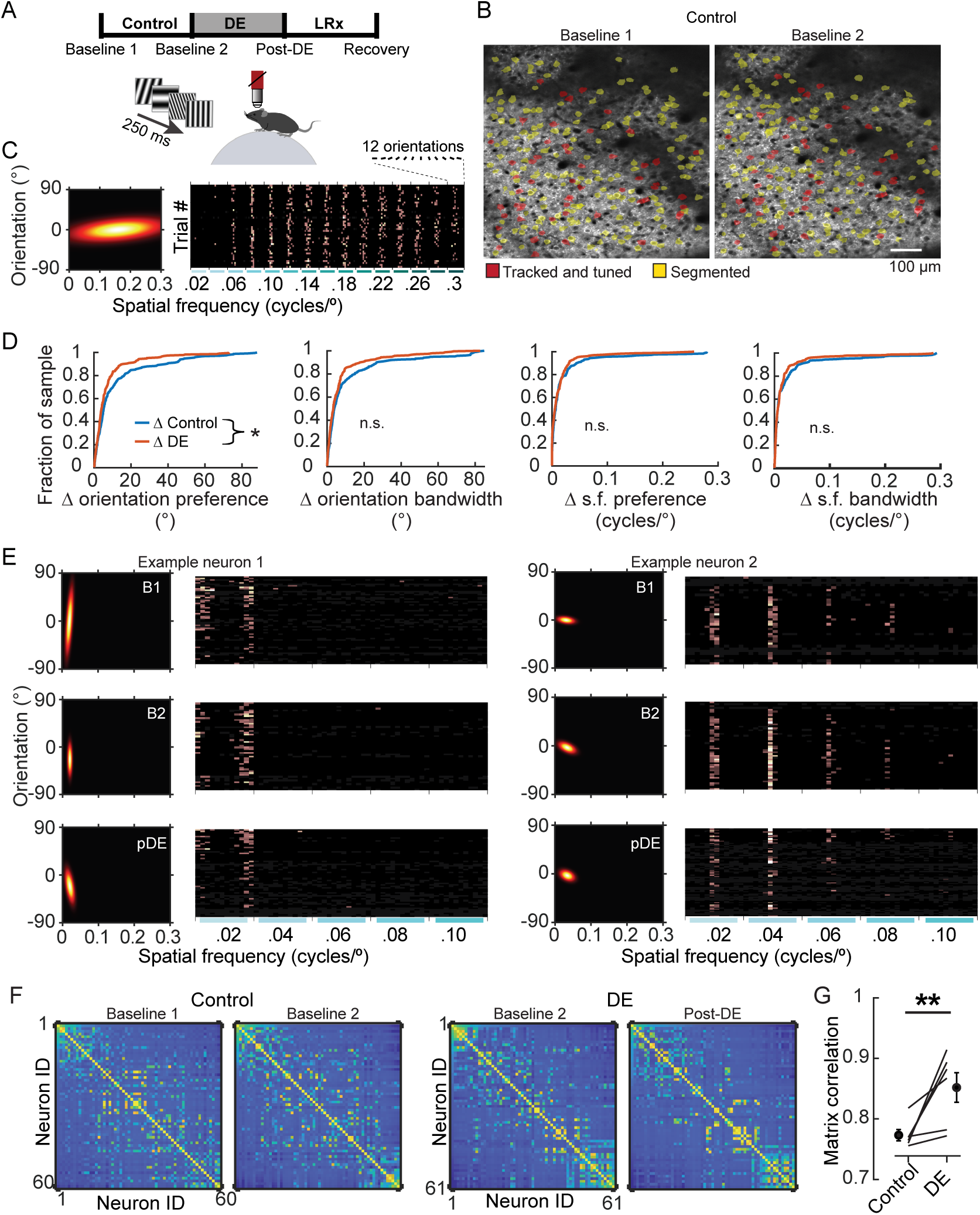
Eight days of dark exposure stabilized orientation preference in V1 neurons. (A) Experimental design; 3 conditions were studied: Control, dark exposure (DE), and LRx, bounded by 4 imaging sessions, as indicated. Imaging sessions were acquired while static gratings were presented to awake, head fixed mice, 8±1 days apart. 12 orientations and 15 spatial frequencies, ranging from 0.02 to 0.3 cycles/° were presented. (B) Example of longitudinal imaging across two sessions. (C) Example neuron tuning curve (left) and trial responses to 180 stimuli (right). Stimulus presentations were sorted post-hoc. (D) The change in four features, as indicated, are plotted for individual neurons, pooled across 6 animals. Orientation preference was significantly less stable (Wilcoxon rank-sum test corrected for 4 multiple comparisons, p= 0.032; approximately 30% of the population is shifted left-word in the DE condition) in the control condition (n= 249 neurons) compared to the DE condition (n=230 neurons). All neurons that were tracked and tuned to grating stimuli on both sessions were included in the analysis. The same set of neurons were used for a given condition, but could differ across conditions, to maximize the number of neurons tracked. (E) Response profiles of two example neurons (tuning curves, left, and trial responses cropped to 0.1 cycles/°, right) are shown for the Baseline 1 (B1), Baseline 2 (B2), and Post-DE (pDE) imaging sessions. (F) Example of signal correlation matrices from one animal comparing the B1 and B2 imaging sessions, and the B2 and Post-DE imaging sessions. (G) Similarity of signal correlation matrices for a given animal (computed as the Pearson’s correlation coefficient between the two signal correlation matrices) was significantly higher (Wilcoxon rank-sum test, p= 0.009) across the DE condition compared to the control condition (n= 6 animals). All tracked and tuned neurons were included, as in ‘D’. * p<0.05, **p<0.01

### Orientation preference of well-tuned neurons is stabilized by dark exposure

Tuning to visual stimuli was assessed by fitting deconvolved responses of neurons that were determined to be responsive to a two-dimensional Gaussian function (see Methods)^29^. The stimulus set consisted of 12 orientations and 15 spatial frequencies spanning a range of 0.02-0.30 cycles/°, resulting in a total of 180 stimuli (**Fig. 1C**). Four parameters were computed from the two-dimensional fits: orientation preference and bandwidth, as well as spatial frequency (s.f.) preference and bandwidth. Stability of each of the four parameters was calculated by comparing the absolute change (Δ) during the control and DE experimental conditions. We took a conservative approach and only considered neurons that were tracked and tuned (i.e. well-fit by the two-dimensional Gaussian function, see Methods for details) on both of the imaging sessions used to calculate the change. DE induced a significant shift in the stability of orientation preference (Wilcoxon rank-sum p= 0.032, **Fig. 1D,E**). The other three parameters trended in the same direction, but the difference between conditions did not reach statistical significance.

Given these trends, it is possible that if all four parameters were considered simultaneously, a clear difference would emerge. To address this possibility, we examined pairwise signal correlation among the neurons that were responsive and tuned (**Fig. 1F**). The similarity of signal correlation in the control condition and in the DE condition was computed for each of the six animals. We found that the similarity was significantly higher in the DE condition (Wilcoxon rank-sum test, p= 0.009; **Fig. 1G**). Thus, in contrast to monocular deprivation^22^, a lack of vision through both eyes did not perturb tuning, and in fact resulted in a net stabilization.

Next we examined whether light reintroduction, which is known to potently induce gene expression even in the adult^30^, disturbed tuning stability of the 4 tuning parameters or pairwise signal correlation (**Fig. 2**). The distribution of the change in orientation preference and bandwidth for the DE condition was indistinguishable from that of the LRx condition (**Fig. 2A**). This was also the case for the other 2 tuning parameters (median Δ s.f. preference, DE: 5.3E-3 and LRx: 4.9E-3; Wilcoxon rank-sum corrected for 4 multiple comparisons, p= 0.999; median Δ s.f. bandwidth, DE: 5.5E-3 and LRx 5.2E-3: Wilcoxon rank-sum corrected for 4 multiple comparisons, p=0.999). Consistent with these results, no difference in the similarity of signal correlation was detected between the DE and LRx conditions. Taken together, these data demonstrate that eight days of DE stabilized tuning response curves to grating stimuli and light reintroduction did not induce a shift in tuning stability when considering those neurons that remained tuned after DE.

**Figure 2.**
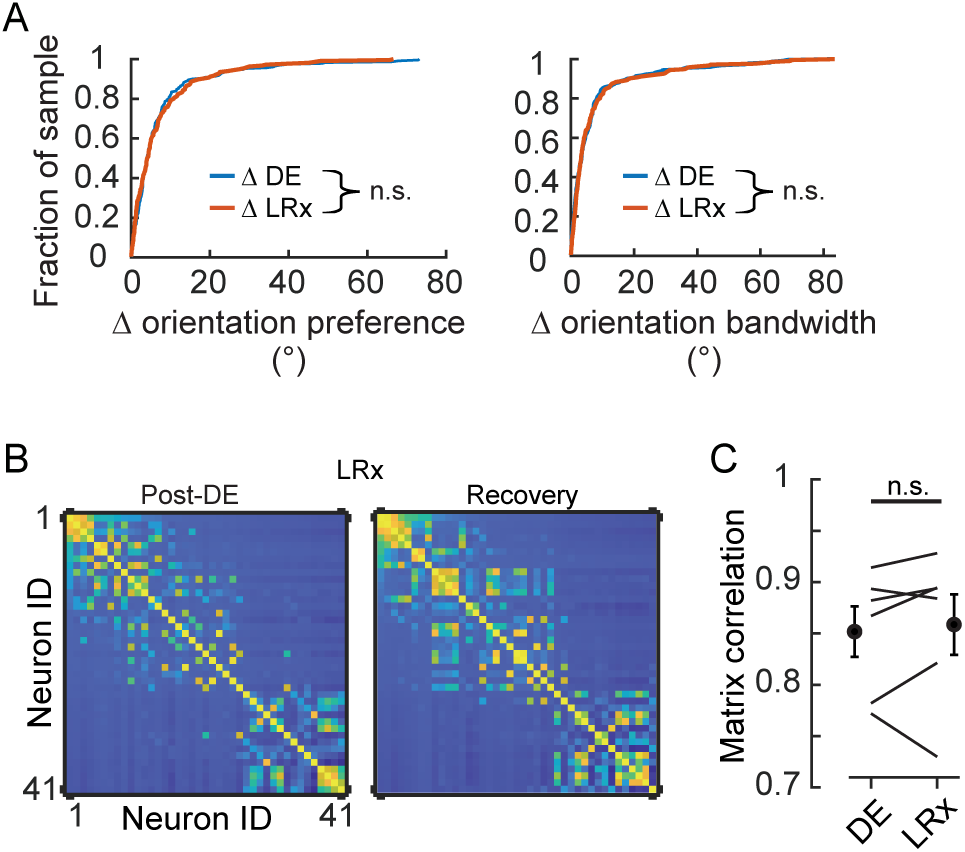
Light reintroduction did not disrupt tuning in V1 neurons. (A) Neither the change in orientation preference or bandwidth was different between DE and LRx conditions (Wilcoxon rank-sum test corrected for 4 multiple comparisons, p=0.795 and p=0.999 respectively; n= 230 and n= 216, respectively). All tracked and tuned neurons were included. (B) Example of signal correlation matrices from one animal comparing the pDE and Rec imaging sessions. (C) Similarity of signal correlation matrices for a given animal (computed as the Pearson’s correlation coefficient) was not different across the DE condition compared to the Recovery condition (Wilcoxon rank-sum test, p= 0.589, n= 6 animals). All tracked and tuned neurons were included.

### Stimulus representation rebounds within eight days of light reintroduction

Not all neurons are well-tuned to grating stimuli, yet such neurons can contribute to visual processing^31^. Therefore, we expanded our analysis to include all tracked neurons, regardless of their responsiveness or tuning characteristics to grating stimuli. We used a k-nearest neighbor (KNN) classifier to decode stimulus identity, as a measure of neural discriminability. First we designed the classifiers such that they were trained separately for each session type, to estimate the amount of stimulus information contained within the network. The identity of the neurons used was the same for all three sessions, referred to as the ‘tracked pool’. Three imaging sessions were examined, the baseline session that immediately preceded dark exposure (Baseline 2, ‘B2’), the session that immediately followed dark exposure (Post-DE, ‘pDE’), and the last session which occurred 8 days after light reintroduction was initiated (Recovery, ‘Rec’). We found that accuracy significantly decreased by 15 ± 3% (±SEM, across animals) on the Post-DE session compared to the Baseline 2 session (paired t-test, p= 0.046), and recovered within eight days (Baseline 2 versus Recovery sessions, paired t-test p= 0.16; **Fig. 3A**). Visual inspection of the confusion matrices indicated that accuracy was degraded across all spatial frequencies (**Fig. 3B,C**). To confirm and quantify this observation, 2 separate classifiers were constructed for all stimuli less than or equal to 0.1 cycles/° (low s.f.) and all stimuli greater than 0.1 cycles/° (high s.f.). In both cases, accuracy was significantly lower on the Post-DE imaging session (paired t-test, p= 0.037 and p= 0.028, respectively; **Fig. 3D**).

**Figure 3.**
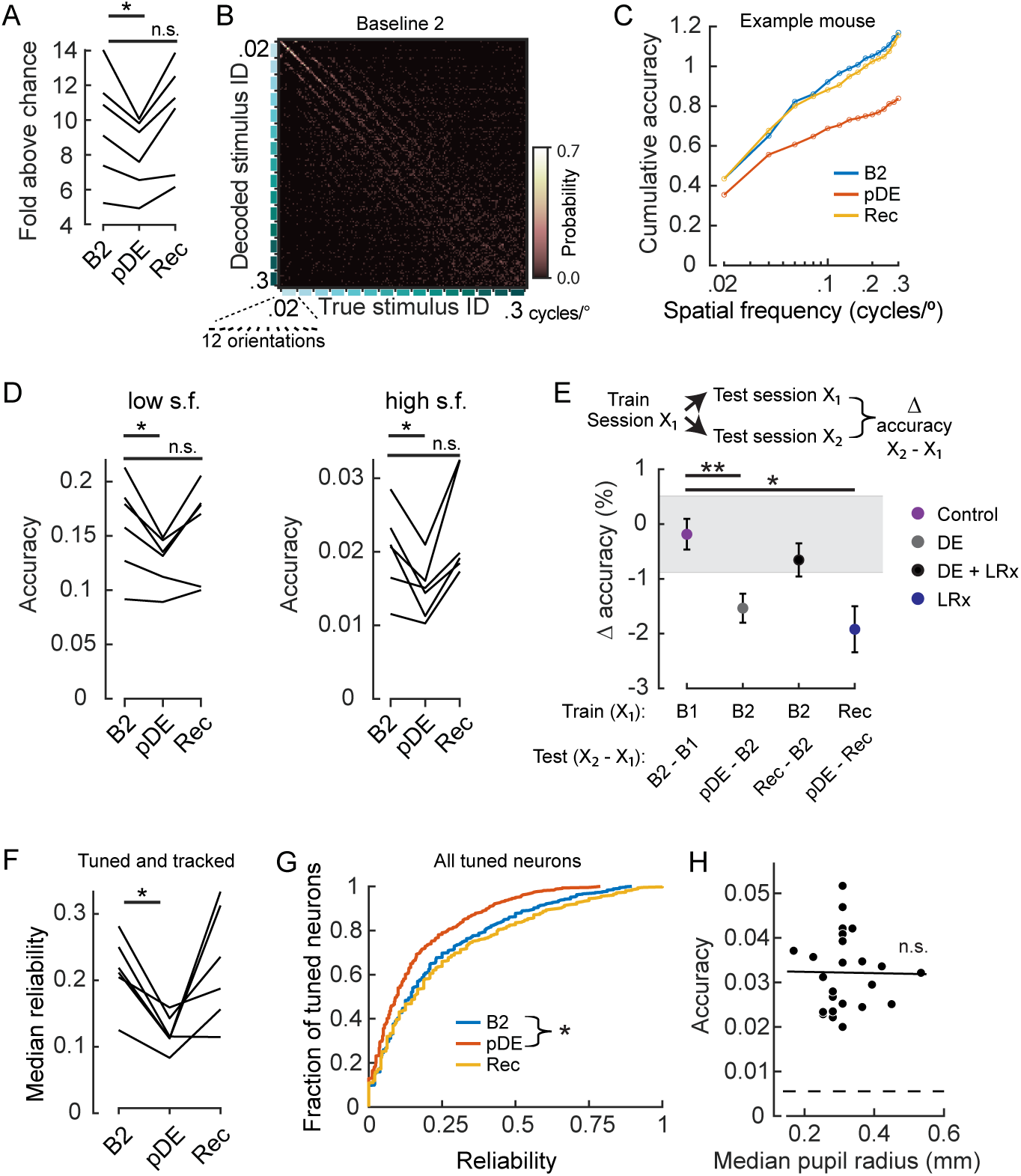
Dark exposure transiently decreased neural discriminability. (A) Classification accuracy was significantly lower (paired t-test corrected for 2 multiple comparisons, p= 0.046) during the Post-DE imaging session (pDE) compared to the Baseline 2 imaging session (B2), and recovered within 8 days of LRx (paired t-test corrected for 2 multiple comparisons, p= 0.16); n= 6 animals. All neurons that were tracked across the B2, pDE and Recovery (Rec) sessions were included. See Methods for number of neurons and trials. Chance was 0.0056. (B) Example of a confusion matrix from one animal during the B2 imaging session. (C) Cumulative sum of classifier accuracy, same mouse as in ‘B’. The classification accuracies of the 12 orientations for a given spatial frequency were averaged to produce 15 data points. (D) Classifiers using only low (≤ 0.1 cycles/°, 60 stimuli) and high (> 0.1 cycles/°, 120 stimuli) spatial frequency stimuli were decoded separately. In both cases, accuracy decreased during the pDE imaging session compared to the B2 session (paired t-test corrected for 2 multiple comparisons, p= 0.037 and p= 0.028, respectively), and recovered within 8 days of LRx (paired t-test corrected for 2 multiple comparisons, p= 0.69 and p= 0.25, respectively). Chance was 0.0167 and 0.0083, respectively. (E) Fixed classifiers were used to quantify representational drift (n= 6 animals, the mean ± SEM). The session used to train the classifier is indicated, as well as the specific sessions tested and the condition label. Note, the rate of drift between the Rec and B2 imaging sessions was comparable to ±1 STD of the baseline drift (gray). In contrast, the rate of drift was significantly higher during the DE and LRx conditions compared to the control condition (paired t-test, corrected for 4 multiple comparisons, ** p= 9E-3 and *p= 0.012). All neurons that were tracked for a given tested session pair were included. (F) The median reliability of responses to the preferred stimulus for a given animal was significantly lower (paired t-test corrected for two multiple comparisons, p= 6E-3) during the pDE imaging session compared to the B2 imaging session; n= 6 animals. All tracked and tuned neurons for a given session were included. (G) Reliability of individual neuron responses to the preferred stimulus, pooled across animals, was significantly reduced (Wilcoxon rank-sum corrected for 2 multiple comparisons, p= 3.E-06) during the pDE imaging session and recovered within 8 days of LRx (Rec). All tuned neurons were included. (H) Decoding accuracy was not correlated with pupil radius (Pearson’s correlation p= 0.95). All imaging sessions from 6 animals were included. Least-squared line indicated. Neuron number (36) was set to the minimum number of tuned neurons among all animals and sessions; trial number (25) was set to the minimum number of trials among all animals and sessions. * p<0.05, **p<0.01

A possible explanation for the decrease in decoding accuracy is that the number of tuned neurons decreased in the pDE session. Such a scenario would also be consistent with the observations reported in Fig. 1D. To determine whether this was the case, the fraction of tuned neurons was compared between the B2 and pDE sessions. Indeed, within the tracked pool, there was a decrease in the number of tuned neurons in the pDE session (paired test corrected for 2 multiple comparisons, p= 0.031; % decrease ± SEM: 11 ± 0.04%; n=6 animals), but not between the B2 and Rec sessions (paired test corrected for 2 multiple comparisons, p= 0.687, 0.01±0.04%). Furthermore, a decrease in response reliability for the preferred stimulus was observed. The median reliability of tuned and tracked neurons for individual mice was significantly lower in the pDE session compared to the B2 session (paired-t test, p= 6E-3; **Fig. 3F**). Similarly, when the reliability of all tuned neurons, regardless of whether tracked, was pooled across mice, the reliability was significantly lower in the pDE session (Wilcoxon rank-sum p= 3E-6, **Fig. 3G**). A potential concern in interpreting these data would be if the pupil was more constricted in the pDE session relative the other sessions. If the pupil diameter was smaller in the pDE session, less light may reach the retina and lead to lower input drive. To determine if this was a factor, we examined pupil radius in relation to the imaging sessions. Rather than a decrease, we detected an increase in pupil radius in the pDE session (median radius in millimeters, B1, B2, pDE, Rec sessions, respectively [± SEM]: 0.29±0.01, 0.28±0.02, 0.41±0.03, and 0.29±0.03). However, pupil diameter did not correlate with decoding accuracy (**Fig. 3H**), therefore we can rule out the possibility that a lower amount of light reaching the retina accounts for the transient increase in representational drift. We noted that on average the mice ran a little more throughout the pDE session (fraction of session spent in locomotion, B1, B2, pDE, Rec sessions, respectively [± SEM]: 0.18±0.05, 0.10±0.05, 0.27±0.06, and 0.07±0.03). Given that pupil diameter is positively correlated with locomotion^32^, it is likely that locomotion drove the increase in pupil diameter.

In our final analysis, to assess the rate of representational drift, fixed classifiers were employed, in which a single session was used to train the classifier. That same session (X_1_) as well as a second session (X_2_), was used for testing the accuracy of stimulus identify classification. The rate of drift was defined as the difference in accuracy between X_2_ and X_1_. The rate of drift was significantly higher during dark exposure (sessions pDE and B2) compared to the control condition (sessions B2 and B1; paired t-test, p= 9E-3; **Fig. 3E**). However, there was not a difference in the change in decoding accuracy across the control condition and between the Rec and the B2 sessions, even though the time span covered was 16 days (DE + LRx) rather than 8 days. These results demonstrate that the stimulus representation not only recovered after transient dark exposure, but that the representation rebounded to its original state. To confirm this interpretation, fixed classifiers were trained using the neural activity from the recovery session and the difference in accuracy between the pDE (the ‘second’ test session in this case) and Rec sessions was computed. As expected, rate of drift was higher than in the control condition (paired t-test, p= 0.012; **Fig. 3E**).

In summary we found that in the adult, stimulus encoding is robust to transient deprivation and is capable of recovering not just in terms of the estimated amount of stimulus information contained in V1, but also that the stimulus representation rebounds to its original form. Light reintroduction, despite initiating a cascade of changes in gene expression^30^, did not persistently disturb stimulus encoding. Thus, using DE as a treatment for amblyopia is not expected to have a negative impact on previously established visual function. Furthermore, our results establish that exposure to naturalistic statistics in the home-cage environment improves neural discriminability, well into adulthood. It will be of interest in future studies to determine whether stimulus discrimination reaches a plateau and requires continuous experience to maintain the plateau, or continues to improve past the classic critical period for ocular dominance plasticity.

### Ideas and Speculation

Our dark exposure experiments revealed that vision has a destabilizing influence on the persistence of orientation preference for approximately 30% of the population of imaged neurons. Similarly, the persistence of pairwise signal correlation among V1 neurons was lower in control conditions compared to the dark exposure condition. These observations are consistent with the theoretical proposal that there is a plastic substrate of neurons with preferentially higher recurrent connectivity that coexist with a stable ‘backbone’ formed by neurons that are resistant to sensory perturbations^33^. Such a functional architecture has the advantage that new information can be integrated into existing networks without perturbing on-going function and could contribute to stable perception while allowing for adaptive flexibility. Also consistent with the proposed functional architecture, we found that that the decoding accuracy of fixed classifiers rebounded to their original, pre-dark exposed state during light reintroduction.

Notably, the development of natural scene processing is protracted relative to grating stimulus encoding in V1^34^. Specifically, a proportion of V1 neurons develop a strong preference for complex scenes relative to simple grating stimuli^34–36^. It will be of future interest to determine whether the development of complex scene processing is facilitated by the presence of neurons that are functionally distinguishable from a stable ‘backbone’, and the extent to which complex-scene preferring neurons emerge from a potentially more plastic population within V1. Such an architecture may be reinforced by a spatial segregation of neuomodulatory input arising from subcortical brain regions associated with reward signals, such as the nucleus basalis ^37–39^. In this scenario, a stable ‘backbone’ could serve the purpose of retaining spatial receptive field position and binocular alignment^40–42^ between the two eyes during the protracted development of complex scene processing.

## Author contributions

BBJ, SMC, and SJK designed the experiments; BBJ and TF collected the data; BBJ, TF, SMC, and SJK analyzed the data, and BBJ and SJK wrote the manuscript.

## Acknowledgements

We thank Jeffrey Good for performing surgeries. Funded by: NIH R01EY024678 (SJK) and The Curci Foundation (SJK and SMC).

## Competing interests

The authors declare that no competing interests exist.

## Methods

### Animal Preparation

All experimental procedures were compliant with the guidelines established by the Institutional Animal Care and Use Committee of Carnegie Mellon University and the National Institutes of Health, and all experimental protocols were approved by the Institutional Animal Care and Use Committee of Carnegie Mellon University. To express the calcium indicator GCaMP6f selectively in excitatory neurons, homozygous Emx1cre mice (Jackson Laboratories, stock number 005628) were crossed with homozygous Ai93/heterozygous Camk2a-tTA mice (Jackson Laboratories, stock number 024108). Experimental mice were heterozygous for all three alleles. Mice were housed in groups of 2-3 per cage, in a 12-hour light/12-hour dark cycle; all imaging sessions started at Zeitgeber time (ZT) 14.5±1, where ZT0 is lights on, and ZT12 is lights off. The same enrichment materials were provided in all cages including a Plexiglas hut and nesting material. See **Table 1** for information on animal sex and genotype. None of the mice used in this study exhibited aberrant, interictal events^34,43^ in V1 or adjacent regions.

Mice (29-31 days old) were anesthetized with isoflurane (3% induction, 1-2% maintenance). A 3-mm diameter craniotomy was made over the primary visual cortex in the left hemisphere, identified by coordinates and landmarks as described in^6^. A stainless-steel bar, used to immobilize the head for recordings, was glued to the right side of the skull and secured with dental cement. The craniotomy was then covered with a double glass assembly in which the diameter of the inner glass was fitted to the craniotomy and sealed with dental cement. Mice were allowed to recover for a minimum of 3 days with ad libitum access to food and water.

### Data Acquisition, Neuron Segmentation, and Neuron Tracking

Two-photon calcium imaging was performed in awake head-fixed mice mounted atop a floating spherical treadmill using a resonant scanning microscope (Neurolabware) outfitted with a 16x Nikon objective (0.80 NA) and 8 kHz resonant scanning mirror. Treadmill motion was recorded using a camera (Dalsa Genie M640-1/3) for off-line analysis of locomotion^29^, and eye blinks were captured using a second camera (Dalsa Genie M1280)^34^. A laser excitation wavelength of 920 nm was used (Coherent, Inc.); green emissions were filtered (Semrock 510/84-50), amplified (Edmund Optics 59-179), and detected with a PMT (Hamamatsu H1 0770B-40). The imaged field of view was 620 × 504 microns, pixel dimensions were 0.85 × 0.98 µm, and the acquisition rate was 15.5 Hz. The acquired image time series were motion-corrected by computing the horizontal and vertical translation of each frame using phase correlation^34^, and individual neurons segmented using the Matlab version of Suite2p toolbox^44^, as described in^34^.

To identify neurons that were tracked across imaging sessions, we registered repeat imaging sessions using the mean intensity image of each session. The mean intensity image for a session was computed by averaging the intensity of each pixel in the aligned calcium image series across time for the entire imaging session (roughly 50000 frames). Then, the mean intensity images of the two sessions were registered using an affine transform with one-plus-one evolutionary optimizer. Once the sessions were registered, the percentage of pixel overlap between the neurons from two sessions was computed. Neurons were accepted to be the same neuron across sessions, if the percentage of overlapping pixels across the two sessions was larger than 75%. On average, there were 160 pixels in a given neuron.

### Visual Stimulation

Static sinusoidal grating stimuli were generated using psychophysics toolbox (http://psychtoolbox.org/) in Matlab (Mathworks, Boston, MA). The stimulus was presented on a screen positioned 25 cm away from the right eye angled at 50° with respect to the midline of the animal. The size of the screen was 64 cm by 40 cm, thereby subtending 142° × 96° of visual angle. The spatial frequency range of the stimulus set was 0.02 cycles/° to 0.3 cycles/° at 0.02 cycles/° interval. The orientations ranged from 0° to 180° at 15° spacing interval, yielding a total of 180 different sinusoidal gratings with 12 different orientations and 15 different spatial frequencies. Each grating was presented for 250 ms consecutively in a random order without interleaved gray screen, this was repeated 4 times and data were saved to disk. This sequence was repeated a minimum of 9 times, resulting in a total of at least 36 trials for a given stimulus. Taking into account trials removed due to locomotion or pupil tracking (see *Quantification of Visual Responses*), a minimum of 25 trials was used in analysis. Two seconds of isolumant gray screen was presented at the onset of each sequence.

### Quantification of Visual Responses

Reverse correlation was used to determine the response window of a given stimulus^45^. The peak in the stimulus-averaged events was observed 194 ms to 320 ms after the stimulus was presented on the screen. Therefore, for each stimulus, the corresponding event activity was computed by averaging the number of events between 194 ms and 320 ms window. We defined this period as the response window for a given stimulus.

A neuron was defined as responsive to visual stimuli when the number of events following a presentation of a visual stimulus was modulated by the stimuli presented. To test for modulation, we performed a one-way analysis of variance (ANOVA, α= 0.01) on the observed events during the response window using stimuli as the factor for each neuron. GCaMP6f expressed in neurons had longer decay than the presentation rate of our stimuli^46^. Therefore, we used deconvolution to remove the effects of decay in calcium fluorescence in quantifying responses of each neuron to our visual stimuli as in^29^. Briefly, amplitude of calcium transients was expressed in units of inferred events. For each segment n, inferred events s_*n*_ were estimated from fluorescence using the following model:

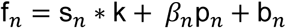

where k is the temporal kernel and b_*n*_ is the baseline fluorescence. Neuropil fluorescence, which is a contamination of the fluorescence signal f_*n*_ from out of focus cell bodies and nearby axons and dendrites, is modeled by p_*n*_, the time course of the neuropil contamination, and, *β*_*n*_ the scaling coefficients. * denotes convolution. Using this model, s_*n*_, k, *β*_*n*_, and b_*n*_ were estimated by a matching pursuit algorithm with L0 constraint, in which spikes were iteratively added and refined until the threshold determined by the variance of the signal was met.

Trials containing locomotion or eye blinks were removed. Pupil location was estimated from eye-tracking videos using a circular Hough Transform algorithm; the algorithm failed to find the pupil on frames during which the mice were blinking. These frames were marked as eye blink frames and removed from further analysis. Trials with locomotion were identified as in^29^. Briefly, after applying a threshold on the luminance intensity of the treadmill motion images, phase correlation was computed between consecutive frames to estimate the translation between the frames. To define a motion threshold, the data were smoothed using a 1s sliding window. Any continuous non-zero movement periods during which the animal’s instantaneous running speed exceeded 10 cm/s threshold for at least one frame were marked as running epochs.

#### Estimation of preferred stimulus and tuning bandwidth

Orientation and spatial frequency preference were determined using a two-dimensional Gaussian model, fit to single trial responses. For neurons that were responsive to grating stimuli, a two-dimensional Gaussian model was fit using nonlinear least-squared regression such that the number of events R as a function of the orientation θ and the spatial frequency φ of the stimulus was

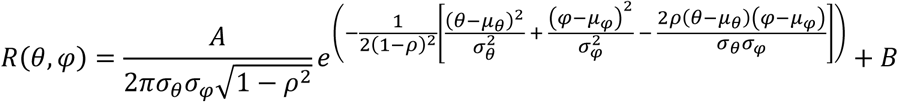

where μ_θ_ was the preferred orientation and μ_φ_ was the preferred spatial frequency of the stimulus, and the σ_θ_ and σ_φ_ described the widths of respective tuning. The covariance of responses for orientation and spatial frequency was captured by the correlation term ρ. A was a parameter accounting for the amplitude of the responses in number of events, while B was the baseline event activity of the cell. For fitting, the lower and the upper bound of allowed values for μ_φ_ was set by the range of the presented stimuli, which was 0.02 to 0.30 cycles/°. The lower bound for σ_θ_ and σ_φ_ was set at 1 degree and 0.001 cycles/° respectively to prevent fits with zero or negative widths. Prior to fitting, the preferred orientation was initialized by estimating the preferred orientation by averaging the response, *R* across all spatial frequencies for a given stimulus orientation, θ and calculating half the complex phase of the value

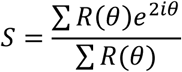

^47,48^. The preferred spatial frequency was initialized by selecting the spatial frequency that generated the maximal significant response at the estimated preferred orientation. For the model above, R^2^ of the fit was used to find neurons with significant tuning. The chance distribution of R^2^ was calculated from fitting the above model with permuted stimulus labels on individual trials 1000 times for each neuron. Neurons whose R^2^ exceeded the 95^th^ percentile of the chance R^2^ distribution were accepted as well-tuned to grating stimuli.

The bandwidths of the Gaussian tuning were described using half-width at half-maximum (HWHM). The HWHM bandwidths for both orientation and spatial frequency were calculated as

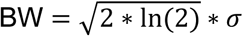

where *σ* was the width parameter of the Gaussian fit.

#### Computation of signal correlation (*Figures 1F,G; 2B*)

Signal correlation ρ^sig^ between a pair of neurons is defined as Pearson’s correlation between the average responses to stimuli^49^. Therefore, we computed pairwise signal correlation between neuron i and neuron j as

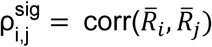

where 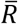 is a vector of average response in number of spikes to 180 sinusoidal gratings for the respective neuron.

#### Stimulus classification (*Figure 3A-E*)

K-nearest-neighbor (KNN) classifiers were used to decode the presented stimuli from vectors of single trial population responses^34,36^. In our case, the k-nearest-neighbor classifier estimated the stimulus identity for a given response vector by identifying the most frequent stimulus identity of its k closest response vectors. To identify the nearest neighbors for a given response vector, we computed the Euclidean distance to the other response vectors. For each session, data was divided so that a single set of response vectors consisted of one trial of each stimulus. This resulted in the number of sets being equal to the number of trials that each stimulus was shown. When the number of trials available was larger than the minimum number of trials, trials were randomly subsampled from the available trials. During decoding, the possible neighbors for a test response vector consisted of all response vectors not belonging to the test set. This ensures an unbiased representation of possible nearest neighbors across stimuli. This process was repeated across each response vector and each set. We reported the performance of this decoding process as accuracy across all response vector tested. Previously we found that a value of k= 4 resulted in the best average rank across mice^50^, therefore we fixed the value of k to 4. Chance performance of the classifiers was 1 divided by the number of stimuli classified. The number of trials and neurons was matched to the minimum number of available trials and neurons across the three sessions, B2, pDE, and Rec. In the case more trials or neurons were available, neuros and trials were randomly subsampled.

To quantify representation drift, we modified the KNN classifier described above. We trained a KNN classifier on the neuronal responses from a single session, and decoded held-out responses from that given session and as well as responses from a second session acquired on a different day. Only the neurons that were tracked in both sessions were used to train and test the classification algorithm. The number of trials was matched to the minimum number of available trials across the two sessions.

The number of neurons and trials used were as follows: Figure 3A, and D, for mouse # 1-6, the number of trials was 42,31,33,27,26,37 for each of the imaging sessions, and the number of neurons was 114,98,75,64,55,96, respectively; Figure 3E, the number of trials ranged between 25 – 44 and the number of neurons ranged between 68 – 167, depending on the animal and condition. Figure 3H, the number of trials was 25 and the number of neurons was 36.

#### Reliability across trials (*Figure 3F,G*)

Reliability was computed as the proportion of trials in which the response amplitude was at least 2 standard deviations above baseline activity, where baseline activity was defined as the activity during presentation of the isolumant gray screen.

### Statistics

Error is reported as standard error of the mean (S.E.M.), unless noted. In the case data were not normally distributed, non-parametric tests were used. Alpha was set to 0.05 unless noted. In the case P values were corrected for multiple comparisons, the number of comparisons is noted in the figure legend; correction was computed using the false discovery rate (FDR) Benjamini-Hochberg procedure.

## Notes

### Competing Interest Statement

The authors have declared no competing interest.

## References

1. Buonomano, D. V & Merzenich, M. M. Cortical plasticity: from synapses to maps. Annu. Rev. Neurosci. 21, 149–186 (1998).

2. Reh, R. K., Dias, B. G., Nelson, C. A. 3rd, Kaufer, D., Werker, J. F., Kolb, B., Levine, J. D. & Hensch, T. K. Critical period regulation across multiple timescales. Proc. Natl. Acad. Sci. U. S. A. 117, 23242–23251 (2020).

3. Hensch, T. K. & Quinlan, E. M. Critical periods in amblyopia. Vis. Neurosci. 35, E014 (2018).

4. Fong, M.-F., Duffy, K. R., Leet, M. P., Candler, C. T. & Bear, M. F. Correction of amblyopia in cats and mice after the critical period. Elife 10, (2021).

5. Jenks, K. R., Kim, T., Pastuzyn, E. D., Okuno, H., Taibi, A. V, Bito, H., Bear, M. F. & Shepherd, J. D. Arc restores juvenile plasticity in adult mouse visual cortex. Proc. Natl. Acad. Sci. U. S. A. 114, 9182–9187 (2017).

6. Feese, B. D., Pafundo, D. E., Schmehl, M. N. & Kuhlman, S. J. Binocular deprivation induces both age-dependent and age-independent forms of plasticity in parvalbumin inhibitory neuron visual response properties. J. Neurophysiol. 119, 738–751 (2018).

7. Murase, S., Winkowski, D., Liu, J., Kanold, P. O. & Quinlan, E. M. Homeostatic regulation of perisynaptic matrix metalloproteinase 9 (MMP9) activity in the amblyopic visual cortex. Elife 8, (2019).

8. Faini, G., Aguirre, A., Landi, S., Lamers, D., Pizzorusso, T., Ratto, G. M., Deleuze, C. & Bacci, A. Perineuronal nets control visual input via thalamic recruitment of cortical PV interneurons. Elife 7, (2018).

9. Jenks, K. R., Tsimring, K., Ip, J. P. K., Zepeda, J. C. & Sur, M. Heterosynaptic Plasticity and the Experience-Dependent Refinement of Developing Neuronal Circuits. Front. Neural Circuits 15, 803401 (2021).

10. Pizzorusso, T., Medini, P., Landi, S., Baldini, S., Berardi, N. & Maffei, L. Structural and functional recovery from early monocular deprivation in adult rats. Proc. Natl. Acad. Sci. U. S. A. 103, 8517–8522 (2006).

11. Pizzorusso, T., Medini, P., Berardi, N., Chierzi, S., Fawcett, J. W. & Maffei, L. Reactivation of ocular dominance plasticity in the adult visual cortex. Science 298, 1248–1251 (2002).

12. Falcone, M. M., Hunter, D. G. & Gaier, E. D. Emerging therapies for amblyopia. Semin. Ophthalmol. 36, 282–288 (2021).

13. Rodríguez, G., Chakraborty, D., Schrode, K. M., Saha, R., Uribe, I., Lauer, A. M. & Lee, H.-K. Cross-Modal Reinstatement of Thalamocortical Plasticity Accelerates Ocular Dominance Plasticity in Adult Mice. Cell Rep. 24, 3433-3440.e4 (2018).

14. Eaton, N. C., Sheehan, H. M. & Quinlan, E. M. Optimization of visual training for full recovery from severe amblyopia in adults. Learn. Mem. 23, 99–103 (2016).

15. He, H.-Y., Hodos, W. & Quinlan, E. M. Visual deprivation reactivates rapid ocular dominance plasticity in adult visual cortex. J. Neurosci. 26, 2951–2955 (2006).

16. Montey, K. L. & Quinlan, E. M. Recovery from chronic monocular deprivation following reactivation of thalamocortical plasticity by dark exposure. Nat. Commun. 2, 317 (2011).

17. Murase, S., Lantz, C. L. & Quinlan, E. M. Light reintroduction after dark exposure reactivates plasticity in adults via perisynaptic activation of MMP-9. Elife 6, (2017).

18. He, H.-Y., Ray, B., Dennis, K. & Quinlan, E. M. Experience-dependent recovery of vision following chronic deprivation amblyopia. Nat. Neurosci. 10, 1134–1136 (2007).

19. Erchova, I., Vasalauskaite, A., Longo, V. & Sengpiel, F. Enhancement of visual cortex plasticity by dark exposure. Philos. Trans. R. Soc. Lond. B. Biol. Sci. 372, (2017).

20. Stodieck, S. K., Greifzu, F., Goetze, B., Schmidt, K.-F. & Lowel, S. Brief dark exposure restored ocular dominance plasticity in aging mice and after a cortical stroke. Exp. Gerontol. 60, 1–11 (2014).

21. Mitchell, D. E., Crowder, N. A. & Duffy, K. R. The critical period for darkness-induced recovery of the vision of the amblyopic eye following early monocular deprivation. J. Vis. 19, 25 (2019).

22. Rose, T., Jaepel, J., Hubener, M. & Bonhoeffer, T. Cell-specific restoration of stimulus preference after monocular deprivation in the visual cortex. Science 352, 1319–1322 (2016).

23. Torrado Pacheco, A., Tilden, E. I., Grutzner, S. M., Lane, B. J., Wu, Y., Hengen, K. B., Gjorgjieva, J. & Turrigiano, G. G. Rapid and active stabilization of visual cortical firing rates across light-dark transitions. Proc. Natl. Acad. Sci. U. S. A. 116, 18068–18077 (2019).

24. Deitch, D., Rubin, A. & Ziv, Y. Representational drift in the mouse visual cortex. Curr. Biol. 31, 4327-4339.e6 (2021).

25. Schoonover, C. E., Ohashi, S. N., Axel, R. & Fink, A. J. P. Representational drift in primary olfactory cortex. Nature 594, 541–546 (2021).

26. Rupasinghe, A., Francis, N., Liu, J., Bowen, Z., Kanold, P. O. & Babadi, B. Direct extraction of signal and noise correlations from two-photon calcium imaging of ensemble neuronal activity. Elife 10, (2021).

27. Margolis, D. J., Lutcke, H., Schulz, K., Haiss, F., Weber, B., Kugler, S., Hasan, M. T. & Helmchen, F. Reorganization of cortical population activity imaged throughout long-term sensory deprivation. Nat. Neurosci. 15, 1539–1546 (2012).

28. McGuire, K. L., Amsalem, O., Sugden, A. U., Ramesh, R. N., Fernando, J., Burgess, C. R. & Andermann, M. L. Visual association cortex links cues with conjunctions of reward and locomotor contexts. Curr. Biol. 32, 1563-1576.e8 (2022).

29. Jeon, B. B., Swain, A. D., Good, J. T., Chase, S. M. & Kuhlman, S. J. Feature selectivity is stable in primary visual cortex across a range of spatial frequencies. Sci. Rep. 8, 15288 (2018).

30. Mardinly, A. R., Spiegel, I., Patrizi, A., Centofante, E., Bazinet, J. E., Tzeng, C. P., Mandel-Brehm, C., Harmin, D. A., Adesnik, H., Fagiolini, M. & Greenberg, M. E. Sensory experience regulates cortical inhibition by inducing IGF1 in VIP neurons. Nature 531, 371–375 (2016).

31. Levy, M., Sporns, O. & MacLean, J. N. Network Analysis of Murine Cortical Dynamics Implicates Untuned Neurons in Visual Stimulus Coding. Cell Rep. 31, 107483 (2020).

32. Reimer, J., Froudarakis, E., Cadwell, C. R., Yatsenko, D., Denfield, G. H. & Tolias, A. S. Pupil fluctuations track fast switching of cortical states during quiet wakefulness. Neuron 84, 355–362 (2014).

33. Sweeney, Y. & Clopath, C. Population coupling predicts the plasticity of stimulus responses in cortical circuits. Elife 9, (2020).

34. Kowalewski, N. N., Kauttonen, J., Stan, P. L., Jeon, B. B., Fuchs, T., Chase, S. M., Lee, T. S. & Kuhlman, S. J. Development of natural scene representation in primary visual cortex requires early postnatal experience. Curr. Biol. 31, 369-380.e5 (2021).

35. Walker, E. Y., Sinz, F. H., Cobos, E., Muhammad, T., Froudarakis, E., Fahey, P. G., Ecker, A. S., Reimer, J., Pitkow, X., Tolias, A. S., de Vries, S. E. J., Lecoq, J. A., Buice, M. A., Groblewski, P. A., Ocker, G. K., Oliver, M., Feng, D., Cain, N., Ledochowitsch, P., Millman, D., Roll, K., Garrett, M., Keenan, T., Kuan, L., Mihalas, S., Olsen, S., Thompson, C., Wakeman, W., Waters, J., Williams, D., Barber, C., Berbesque, N., Blanchard, B., Bowles, N., Caldejon, S. D., Casal, L., Cho, A., Cross, S., Dang, C., Dolbeare, T., Edwards, M., Galbraith, J., Gaudreault, N., Gilbert, T. L., Griffin, F., Hargrave, P., Howard, R., Huang, L., Jewell, S., Keller, N., Knoblich, U., Larkin, J. D., Larsen, R., Lau, C., Lee, E., Lee, F., Leon, A., Li, L., Long, F., Luviano, J., Mace, K., Nguyen, T., Perkins, J., Robertson, M., Seid, S., Shea-Brown, E., Shi, J., Sjoquist, N., Slaughterbeck, C., Sullivan, D., Valenza, R., White, C., Williford, A., Witten, D. M., Zhuang, J., Zeng, H., Farrell, C., Ng, L., Bernard, A., Phillips, J. W., Reid, R. C. & Koch, C. Inception loops discover what excites neurons most using deep predictive models. Nat. Neurosci. 23, 138–151 (2020).

36. de Vries, S. E. J., Lecoq, J. A., Buice, M. A., Groblewski, P. A., Ocker, G. K., Oliver, M., Feng, D., Cain, N., Ledochowitsch, P., Millman, D., Roll, K., Garrett, M., Keenan, T., Kuan, L., Mihalas, S., Olsen, S., Thompson, C., Wakeman, W., Waters, J., Williams, D., Barber, C., Berbesque, N., Blanchard, B., Bowles, N., Caldejon, S. D., Casal, L., Cho, A., Cross, S., Dang, C., Dolbeare, T., Edwards, M., Galbraith, J., Gaudreault, N., Gilbert, T. L., Griffin, F., Hargrave, P., Howard, R., Huang, L., Jewell, S., Keller, N., Knoblich, U., Larkin, J. D., Larsen, R., Lau, C., Lee, E., Lee, F., Leon, A., Li, L., Long, F., Luviano, J., Mace, K., Nguyen, T., Perkins, J., Robertson, M., Seid, S., Shea-Brown, E., Shi, J., Sjoquist, N., Slaughterbeck, C., Sullivan, D., Valenza, R., White, C., Williford, A., Witten, D. M., Zhuang, J., Zeng, H., Farrell, C., Ng, L., Bernard, A., Phillips, J. W., Reid, R. C. & Koch, C. A large-scale standardized physiological survey reveals functional organization of the mouse visual cortex. Nat. Neurosci. 23, 138–151 (2020).

37. Pafundo, D. E., Nicholas, M. A., Zhang, R. & Kuhlman, S. J. Top-Down-Mediated Facilitation in the Visual Cortex Is Gated by Subcortical Neuromodulation. J. Neurosci. 36, 2904–2914 (2016).

38. Hangya, B., Ranade, S. P., Lorenc, M. & Kepecs, A. Central Cholinergic Neurons Are Rapidly Recruited by Reinforcement Feedback. Cell 162, 1155–1168 (2015).

39. Ji, W., Gamanut, R., Bista, P., D’Souza, R. D., Wang, Q. & Burkhalter, A. Modularity in the Organization of Mouse Primary Visual Cortex. Neuron 87, 632–643 (2015).

40. Wang, B.-S., Sarnaik, R. & Cang, J. Critical period plasticity matches binocular orientation preference in the visual cortex. Neuron 65, 246–256 (2010).

41. Sarnaik, R.,Wang, B.-S. & Cang, J. Experience-dependent and independent binocular correspondence of receptive field subregions in mouse visual cortex. Cereb. Cortex 24, 1658–1670 (2014).

42. Chang, J. T., Whitney, D. & Fitzpatrick, D. Experience-Dependent Reorganization Drives Development of a Binocularly Unified Cortical Representation of Orientation. Neuron 107, 338-350.e5 (2020).

43. Steinmetz, N. A., Buetfering, C., Lecoq, J., Lee, C. R., Peters, A. J., Jacobs, E. A. K., Coen, P., Ollerenshaw, D. R., Valley, M. T., de Vries, S. E. J., Garrett, M., Zhuang, J., Groblewski, P. A., Manavi, S., Miles, J., White, C., Lee, E., Griffin, F., Larkin, J. D., Roll, K., Cross, S., Nguyen, T. V, Larsen, R., Pendergraft, J., Daigle, T., Tasic, B., Thompson, C. L., Waters, J., Olsen, S., Margolis, D. J., Zeng, H., Hausser, M., Carandini, M. & Harris, K. D. Aberrant cortical activity in multiple GCaMP6-expressing transgenic mouse lines. eNeuro 4, (2017).

44. Pachitariu, M., Stringer, C., Dipoppa, M., Schröder, S., Rossi, L. F., Dalgleish, H., Carandini, M. & Harris, K. D. Suite2p: beyond 10,000 neurons with standard two-photon microscopy. bioRxiv (2017). doi:10.1101/061507

45. Ringach, D. L., Sapiro, G. & Shapley, R. A subspace reverse-correlation technique for the study of visual neurons. Vision Res. 37, 2455–2464 (1997).

46. Chen, T.-W., Wardill, T. J., Sun, Y., Pulver, S. R., Renninger, S. L., Baohan, A., Schreiter, E. R., Kerr, R. A., Orger, M. B., Jayaraman, V., Looger, L. L., Svoboda, K. & Kim, D. S. Ultrasensitive fluorescent proteins for imaging neuronal activity. Nature 499, 295–300 (2013).

47. Niell, C. M. & Stryker, M. P. Highly selective receptive fields in mouse visual cortex. J. Neurosci. 28, 7520–7536 (2008).

48. Kuhlman, S. J., Tring, E. & Trachtenberg, J. T. Fast-spiking interneurons have an initial orientation bias that is lost with vision. Nat. Neurosci. 14, 1121–1123 (2011).

49. Averbeck, B. B., Latham, P. E. & Pouget, A. Neural correlations, population coding and 1 computation. Nat. Rev. Neurosci. 7, 358–366 (2006).

50. Jeon, B. B., Chase, S. M. & Kuhlman, S. J. New skill acquisition does not perturb existing function in sensory cortex. bioRxiv (2021). doi:10.1101/2021.02.08.430302

